# Spatial proteomic mapping of the human and mouse retina using IBEX

**DOI:** 10.64898/2025.12.19.695231

**Authors:** Yuxuan Meng, Jakub Kubiak, Zuzanna Dzieniak, Lorna Fowler, Rose Avient, Jason Hopley, Linyulong Li, Chaoran Li, Yuan Tian, Bruno Charbit, Colin J. Chu

**Author notes:** These authors contributed equally to this work (Yuxuan Meng, Jakub Kubiak).

## Abstract

**Purpose:** To generate a comparative spatial proteomic atlas of the human and mouse retina using a refined highly multiplexed immunohistochemistry technique called Iterative Bleaching Extends Multiplexity (IBEX).

**Methods:** We refined the IBEX workflow by integrating an antibody removal option alongside chemical bleaching. This dual strategy enabled removal of the entire antibody complex allowing for the flexible use of antibodies from the same host species across iterative cycles. Spatial proteomic data were cross-correlated to existing scRNAseq datasets. We also coupled this workflow with super-resolution imaging via deconvolution and applied it to the retina of healthy human, mouse and the Crb1^*rd8*^ mouse model.

**Results:** We successfully imaged over 25 protein markers on human and mouse tissue sections, to generate spatial atlases of the major retinal cell populations. Cross-species protein expression was compared to scRNAseq datasets to identify protein and transcript disparities. Super-resolution IBEX delineated the ultrastructural features of the outer limiting membrane (OLM), identifying CD44 as a core structural component tightly co-localised with a highly organised F-actin belt within Müller glial endfeet. Using the Crb1^*rd8*^ mouse model, disruption of this complex was spatially associated with rosette formation and OLM structural failure.

**Conclusion:** Spatial proteomic atlases of the human and mouse retina were generated and correlated to transcriptional data, revealing insights into the arrangement of major retinal cell populations and OLM structure.

## 1 Introduction

The retina is the highly organised and metabolically active neural tissue within the eye responsible for the detection of light.[1, 2] Its function relies on the precise spatial arrangement and intricate interplay between its neuronal, macroglial, microglial, and vascular components.[3] While single-cell RNA sequencing (scRNAseq) revolutionised our understanding of cellular heterogeneity[4, 5], the required tissue dissociation inherently sacrifices crucial spatial information. Conversely, traditional immunohistochemistry (IHC) is typically limited to the simultaneous detection of 3-5 protein markers that fails to capture the complexity of the cellular environment and signalling networks.

To address these limitations, spatial proteomic technologies have emerged as powerful tools named Nature Method of the Year in 2024. This included Iterative Bleaching Extends Multiplexity (IBEX) which has enabled the profiling of follicular lymphoma to thymic tissue.[6–10] It is an established open-source technique that uses iterative cycles of antibody labelling, imaging, and chemical bleaching to visualise dozens of protein markers on a single tissue section at high resolution while preserving tissue architecture.

Despite its power, the standard IBEX protocol faces challenges in flexibility. The technique relies heavily on commercially available antibodies directly conjugated to fluorophores, which can limit the availability of targets especially in underexplored tissues such as the retina. Furthermore, the standard chemical bleaching (e.g., using lithium borohydride, LiBH_4_) only quenches fluorophores without removing the antibody complex. The choice of available commercial antibodies is often limited due to the host species used to generate both unconjugated and conjugated primary antibodies. Due to secondary antibody cross-reactivity, this usually prevents the use of multiple unconjugated primary antibodies from the same host species (e.g., rabbit or mouse) over different imaging cycles, severely restricting panel design.

This study addressed this challenge by establishing and validating an enhanced IBEX protocol to generate spatial proteomic maps of the human and mouse retina to enable wider discovery research and a deeper characterisation of the retina. Our strategy integrated an antibody removal reagent as an alternative to chemical bleaching. This modification allowed for flexible use of the vast library of well-validated unconjugated retinal antibodies, dramatically expanding the capacity for multiplexing. We utilised this optimised platform to generate spatial proteomic maps of the healthy retina and compared them with existing transcriptomic data. Finally, we integrated super-resolution microscopy with IBEX to investigate the pathophysiology of the outer limiting membrane (OLM) in healthy human, C57BL/6J wildtype mouse and the Crb1^*rd8*^ mouse model of inherited retinal degeneration.

## 2 Results

### 2.1 A refined IBEX workflow enables generation of a spatial proteomic atlas of the healthy human retina

To generate a spatial protein atlas of the human retina using highly multiplexed immunohistochemistry, we refined the IBEX method to build on our published work.[11, 12] The existing protocol relies on chemical inactivation of fluorophores with LiBH_4_, but we now demonstrate flexible integration of a commercial antibody removal reagent into IBEX to also strip bound antibody complexes where required (**Figure 1A**). It overcame a core limitation of the standard IBEX protocol: the inability to use multiple unconjugated primary antibodies from the same host species (e.g., rabbit polyclonals) over different imaging cycles due to the cross-reactivity of secondary antibodies. In tissues such as the retina, for which dedicated commercial platforms or reagent panels do not readily exist, this IBEX modification facilitates greater flexibility in creating novel panels, and more accessible use of existing libraries of retina validated primary antibodies. Furthermore, expanded fluorophore options become available as using Alexa Fluor 594, previously confirmed as resistant to LiBH_4_ bleaching, is now viable (**Supplementary Figure S1**).[11]

**Figure 1.**
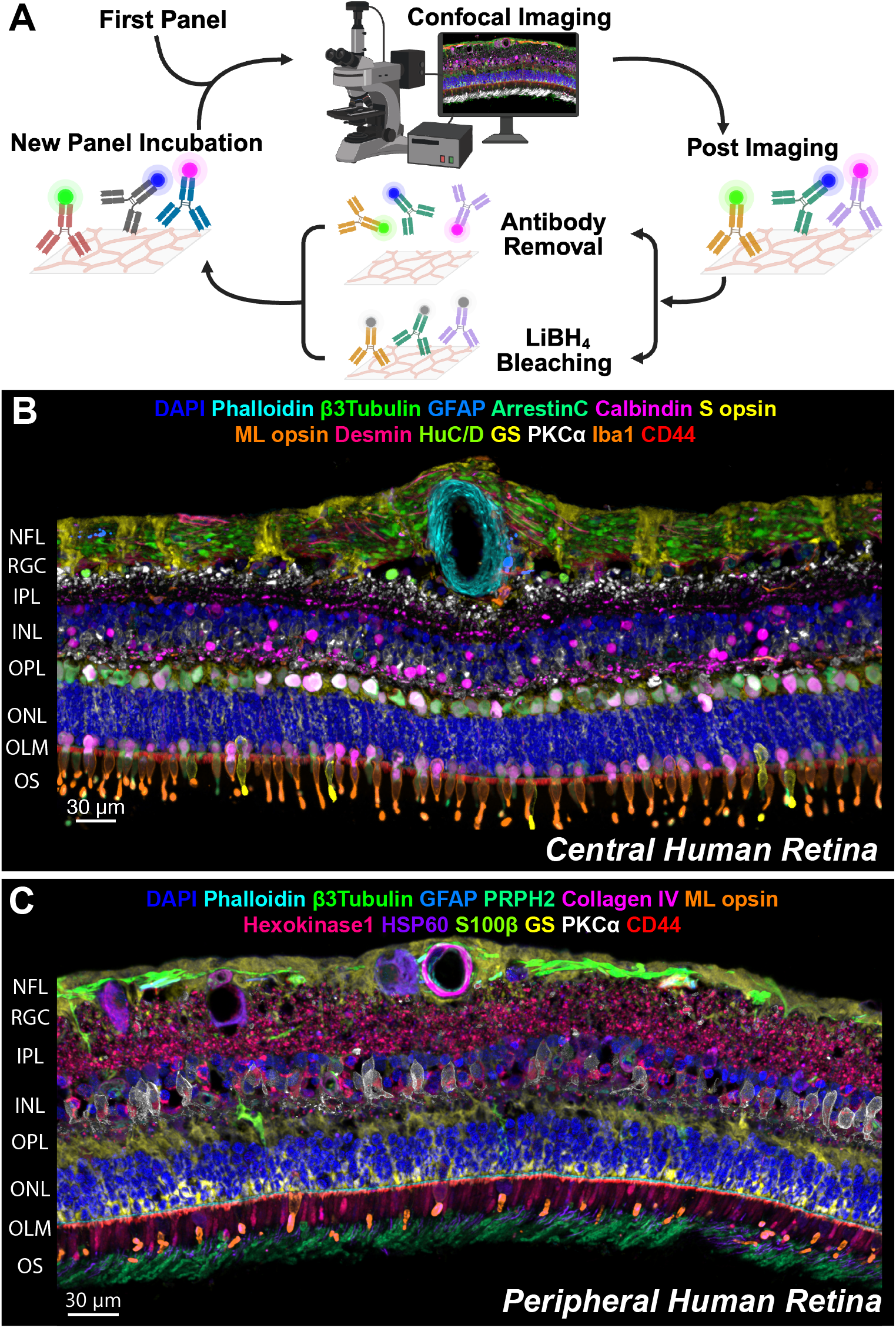
A refined IBEX protocol facilitates generation of a spatial proteomic atlas of the healthy human retina. **(A)** Schematic of the refined IBEX protocol with the inclusion of an antibody complex-removal option. **(B)** Confocal images of healthy central human retina as visualised using the refined IBEX protocol (13 out of 23 parameters displayed). **(C)** Confocal images of a healthy peripheral human retina (13 out of 29 parameters displayed). The representative composites of the human retina show nuclei counterstained with DAPI and markers visualising the retinal cell types spanning neuronal, glial, microglial and vascular compartments.

Using this refined workflow, we generated an atlas of the human retina by imaging over 25 protein markers on fixed frozen sections (See **Supplementary Table S1** for human-reactive antibodies), to cover the major retinal cell types in both central and peripheral regions of healthy human retina (**Figure 1B and 1C**). We confirmed that antibody removal should be done in early panels as after four consecutive rounds it can affect epitope stability.

IBEX allowed precise simultaneous spatial visualisation of the major cell types in the retina (**Figure 2A**) and delineated its fine laminar organization including neuronal structures such as ganglion cells (via *β*3-tubulin or HuC/D), bipolar cells (PKC*α* or CHX10), amacrine or horizontal cells (Calretinin, Calbindin) and photoreceptors (ARR-C for cones and rhodopsin for rods).

**Figure 2.**
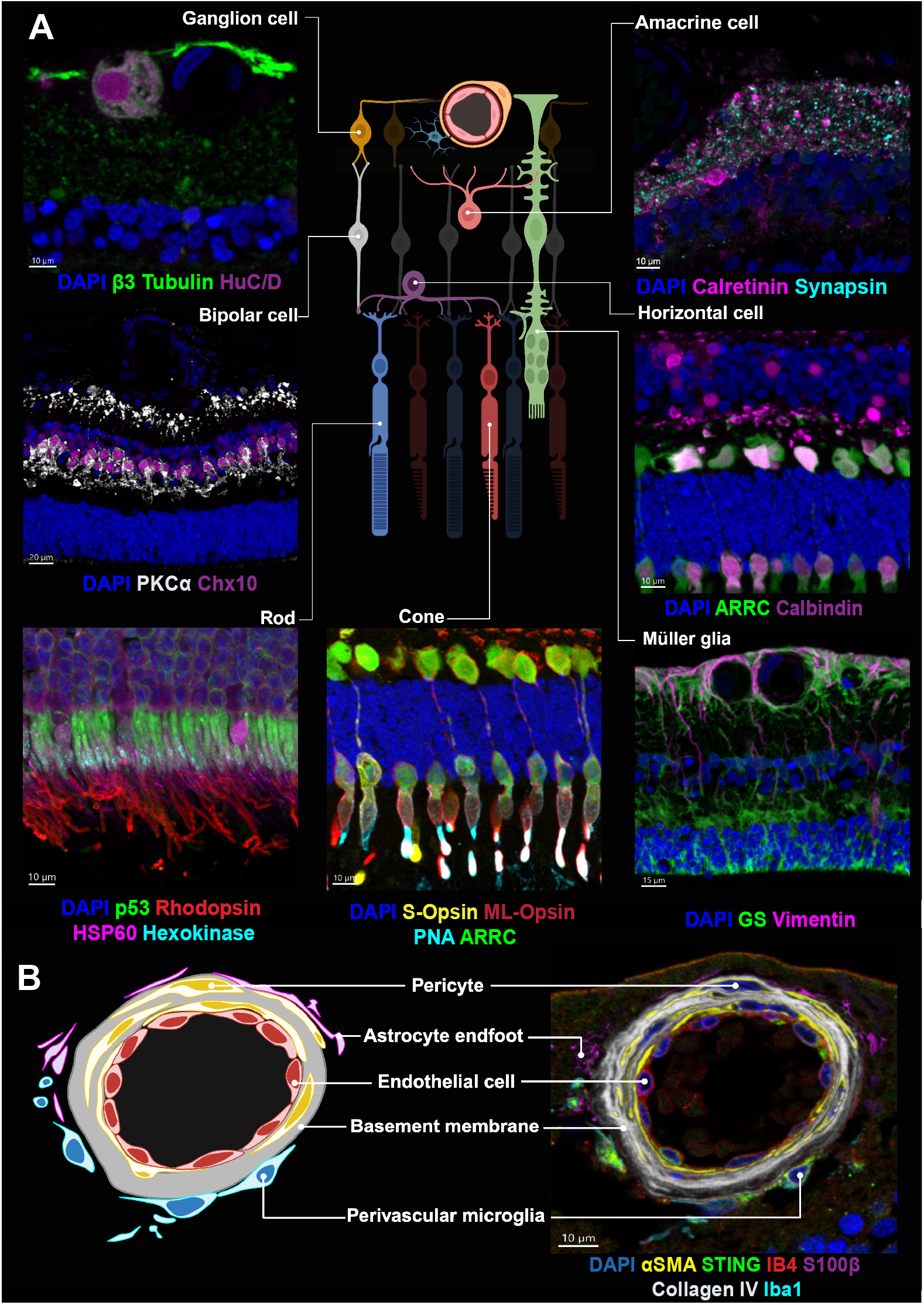
IBEX mapping of healthy human retina identifies major cell types and their respective spatial interactions. **(A)** Major cell types of the human retina displayed with relevant overlapping protein markers. All images were taken from the same tissue. **(B)** IBEX can be used to subcategorise anatomical structures such as the perivascular unit.

Although many retinal antibody markers have been individually published, they have not been imaged in combination on the same tissue section except with 3-5 markers IHC, which does not allow for the simultaneous visualisation of all key structural retinal cells to examine global changes and cellular interactions, which become of greater importance in heterogenous disease states.

The neurovascular unit is a specific example of a complex retinal structure that requires multiple protein markers simultaneously to characterise, which can be achieved using IBEX (**Figure 2B)**. To delineate the multilayered structure and interacting cells of the retinal blood vessel walls we used isolectin B4 (IB4) to label vascular endothelial cells, Collagen IV (COL IV) to outline the basement membrane, *α*-smooth muscle actin (*α*-SMA) to mark pericytes and smooth muscle cells of the vessel wall, and IBA1 to identify perivascular macrophages.

### 2.2 The spatial proteomic atlas of the mouse retina incorporates cross-reactive human-reactive protein markers

Using refined IBEX, we also generated a cellular atlas of the mouse retina including over 20 protein markers on fixed frozen sections. (See **Supplementary Table S2** for mouse-reactive antibodies). Critically, we generated this atlas using 15 of the same human-reactive antibodies which exhibited similar or identical staining pattern in the mouse (**Figure 3A**). Mouse retina features were consistent with the matched human retinal layers across similar physical structures and cellular subtypes (**Figure 3B**).

**Figure 3.**
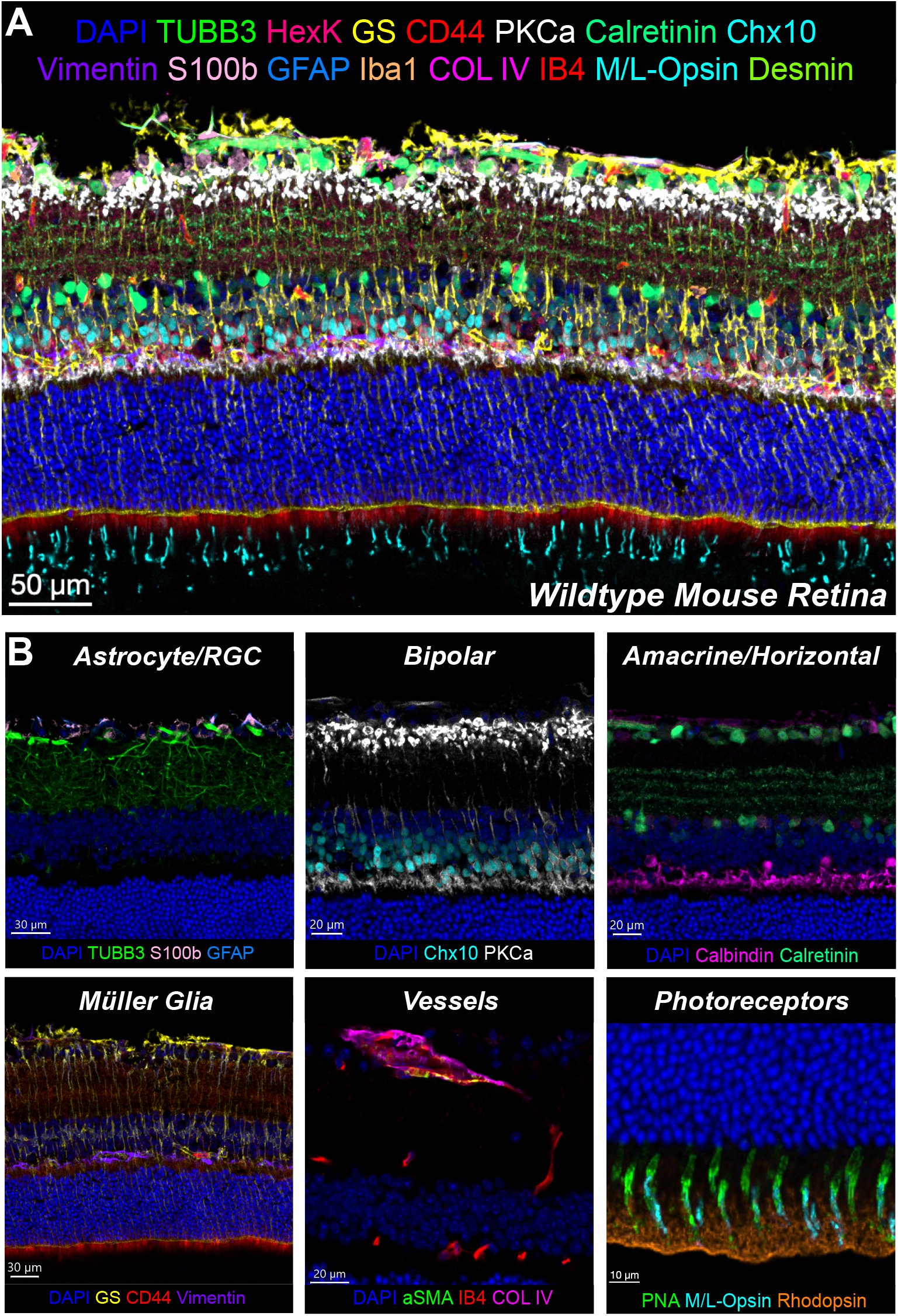
Spatial proteomic cell atlas of the mouse retina. **(A)** Confocal images of a representative healthy mouse retina generated using IBEX. 16 of 28 markers shown for clarity. **(B)** Magnified insets of major cell types and structures of the mouse retina displayed with relevant overlapping protein markers.

To further interpret the findings, we reconciled it with published single-cell RNA sequencing datasets for both mouse and human retinas. **Figure 4A** compares key cross-species reactive proteins and their corresponding transcriptional expression profiles. Mouse and human RNA transcripts and their respective proteins had broadly consistent locations of expression between species (such as IBA1, GFAP or TUBB3, **Figure 4B**).

**Figure 4.**
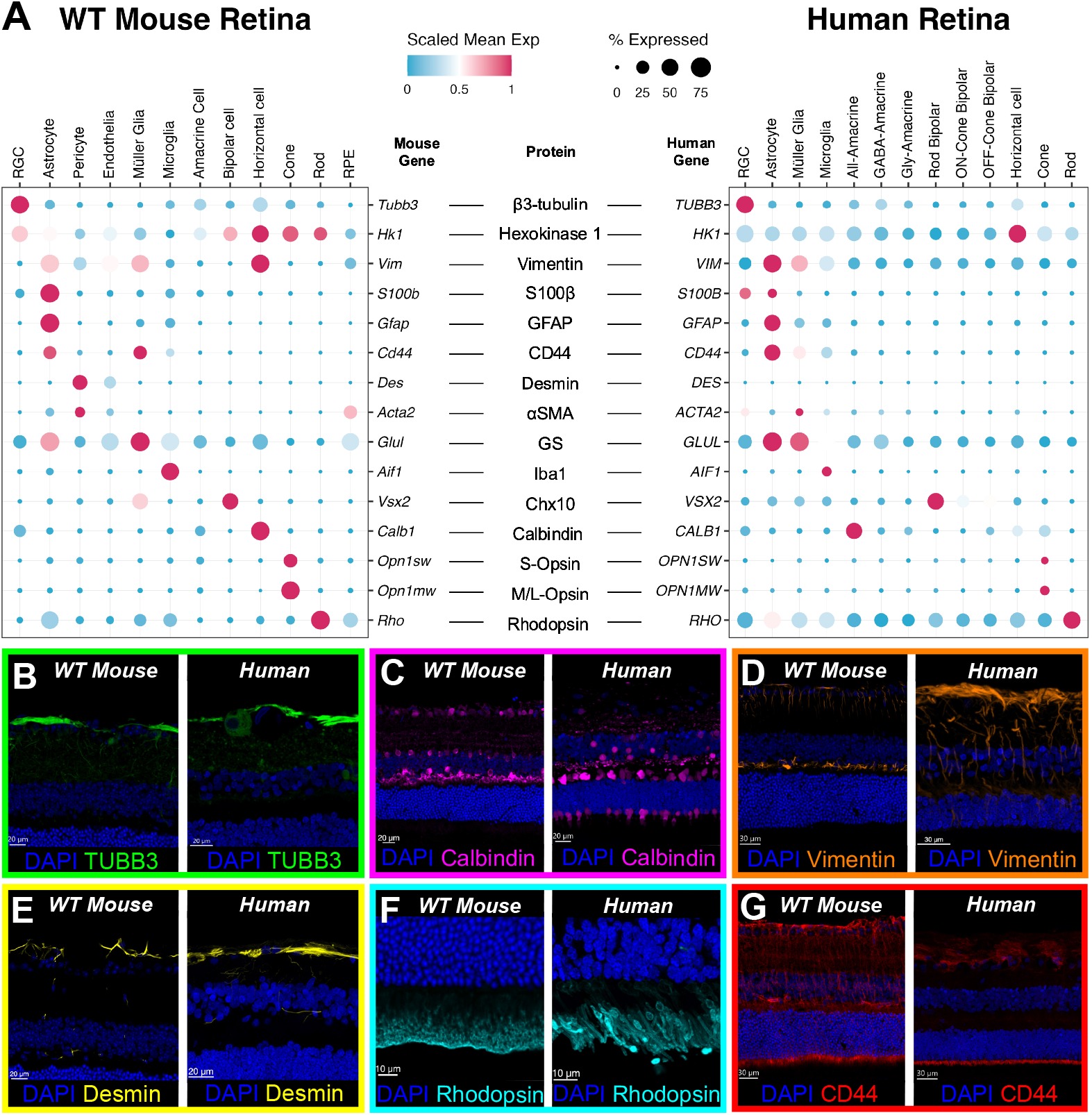
Cross-species comparison of human and mouse retina protein atlases highlights biological disparities with transcriptomic-level data. **(A)** Dot plot visualisation of scRNAseq data for mouse and human retina reconciled against cross-species reactive antibody target proteins. **(B-D)** TUBB3, Calbindin, and Vimentin immunostaining align with their respective transcriptomic profiles in both species. Calbindin exhibited cone staining in humans that was not observed in mouse retina. Vimentin exhibited strong staining at the mouse retina outer plexiform layer, corresponding to horizontal cell staining. **(E)** Desmin protein expression was similar in both species counter to human scRNAseq. **(F)** Rhodopsin protein was uniquely detected in the outer segments only, while RNA expression was ubiquitous. **(G)** Strong CD44 protein detection was localised at the OLM which would not be determined using transcriptomic data.

Disparities between RNA and protein across species were noted. For example, calbindin, as a canonical marker of horizontal cells, also exhibited transcriptional and protein expression within cone photoreceptors in the human retina (**Supplementary Figure S2**), but appears unexpressed within the mouse. **Figure 4C** shows this with the human retina exhibiting calbindin staining at the outer nuclear layer (ONL) and inner segments, which is absent in the mouse.

Vimentin, classically considered as a unique retinal macroglial marker, has a transcriptomic profile demonstrating high expression in horizontal cells in mice. IBEX staining was consistent with this revealing vimentin protein staining at the outer plexiform layer, a feature that is absent in human tissue **(Figure 4D)**. Colocalization within the same IBEX sample with the horizontal cell marker, calbindin, confirmed vimentin co-staining in the mouse (**Supplementary Figure S3**).

Desmin was not observed in any clusters within the human retina transcriptomic datasets but protein staining consistently revealed desmin localisation comparable to the mouse retina, as a distinct feature around the nerve fibre layer (**Figure 4E**). Rhodopsin protein was only exhibited in the outer segments of photoreceptors (**Figure 4F**), in contrast to its transcriptomic profile indicating a widespread rhodopsin expression in all clusters. This is a recognised technical artefact in scRNAseq, as the structural fragility of the photoreceptors makes them sensitive to dissociation methods, potentially cross-contaminating droplets and so confounding cell clustering.[13]

Finally, we observed CD44 as a marker in both mouse and human retina that was transcriptionally expressed on astrocytes and Müller glia. IBEX staining (**Figure 4G**) shows this to be distributed on opposing sides of the retina, with the end-feet of the Müller glia cells exhibiting concentration of CD44 ciliation near to the outer limiting membrane. This is a distinction that could not be elucidated from transcriptomics alone, and is a structure that is underexplored in the literature.[14–18]

### 2.3 Super-resolution IBEX imaging by LIGHTNING deconvolution characterises Outer Limiting Membrane composition in human and mouse retina

We adapted the IBEX technique to integrate semi super-resolution confocal microscopy via Leica LIGHTNING deconvolution to characterise the composition of the OLM boundary with a particular focus on CD44. The OLM boundary is a laminated structure defined by the adherens junctions between Müller glia and photoreceptor cells, observed in both healthy human and wildtype mouse retina (**Figure 5A** and **5B** respectively). We specifically examined the Crb1^*rd8*^ mouse model of inherited retinal degeneration as it is strongly associated with OLM disruption via adhesion changes in the CRB1 complex.[19]

**Figure 5.**
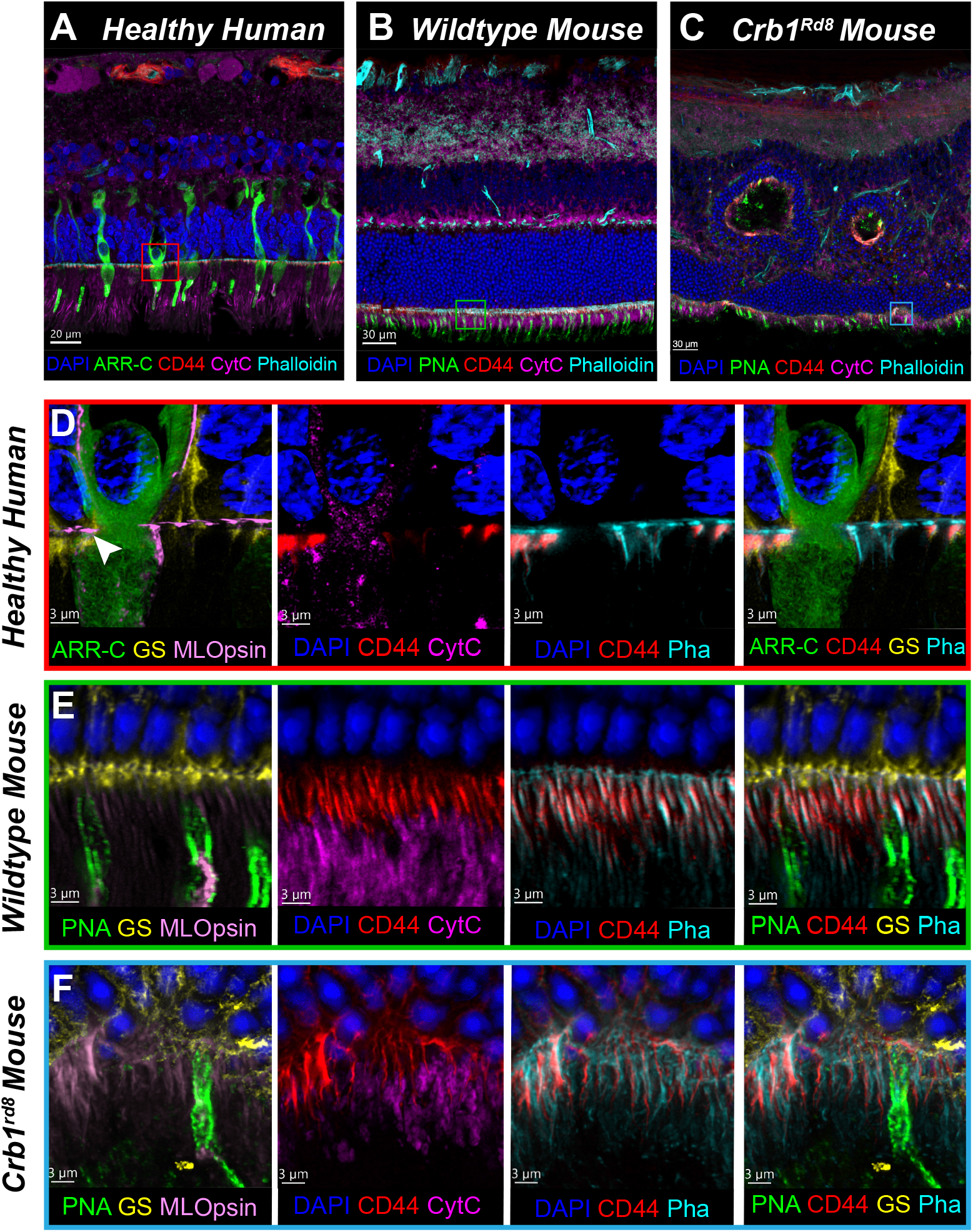
Super-resolution IBEX deconvolves the outer limiting membrane retinal structure. IBEX imaging of a healthy human retina, wildtype, and Crb1^rd8^ mouse retina (**A-C** respectively) visualised at standard resolution. The boxes display the areas of super-resolution imaging. **(D)** Super-resolution image of healthy human retina at the OLM region including a single cone photoreceptor. White arrow indicates the GS-Phalloidin-CD44 interface. **(E)** Wildtype mouse retina exhibited a similar OLM interface pattern, while **(F)** Crb1^rd8^ retina, imaged underneath a rosette, has disrupted OLM structures with Muller glia retraction. Scale Bars: **(A)** 20μm **(B-C)** 30μm **(D-F)** 3 μm.

In the Crb1^*rd8*^ mouse retina, displaced photoreceptor nuclei known as rosettes are formed in the degenerative process (**Figure 5C)**, but the dynamics of their formation and composition are incompletely characterised.[20, 21] Using IBEX, we observe that rosettes are most likely invaginations occurring at the OLM and drawing photoreceptors upwards towards the inner nuclear layer – we note the presence of both rod and cone proteins within the rosettes. These rosettes also contained CD44 and phalloidin, providing evidence that this may be a complete invagination incorporating the OLM, and not just a focal folding or degeneration of the ONL.

Using super-resolution imaging, we could compare the structural configuration of the OLM boundary between human, wildtype mouse and Crb1^*rd8*^ mouse retina. Healthy human and mouse retinas revealed a previously undescribed structural feature: a distinct linear “gap” within the Müller glial apical endfeet (marked by GS) that was intensely stained by phalloidin (indicated by arrow in **Figure 5D** and **Supplementary Figure S4)**. CD44 was further found to be ciliated underneath the phalloidin band and spatially above mitochondria (marked by Cytochrome C) and cone-specific proteins (Arrestin-C, M/L-opsin). Comparative visualisation with the mouse retina (**Figure 5E**) showcased a similar GS-Phalloidin-CD44 interface, albeit the distinct “gap” structure observed in the human retina was not seen in the mouse.

The Crb1^*rd8*^ model at 28 weeks (**Figure 5F**) manifested a clear breakdown of the GS-Phalloidin-CD44 interface with Muller glia endfeet retraction, suggesting indirect spatiotemporal evidence that rosettes may be formed from the OLM pulling into the ONL. With multiplexed imaging, we observed a lower density of PNA-positive cone photoreceptors underneath the rosettes and disruption of inner segment mitochondrial processes aligning with the literature.[21]

## 3 Discussion

Transcriptomics has proven an important resource for the scientific community to define the cellular subtypes of the retina and individual cell state profiles. While there are increasing numbers of commercial highly multiplexed immunohistochemistry platforms, such as CODEX, CycIF, or Imaging Mass Cytometry, these typically feature antibody panels optimised for common tissue types and research areas, and have not included the retina. We now describe the use of antibody removal reagents as a refinement to the flexible and open-source IBEX protocol which was used to generate spatial proteomic atlases of the human and mouse retina and correlated cross-reactive proteins with their respective gene transcript levels acquired from previous scRNAseq datasets. Furthermore, we integrated a deconvolution super-resolution approach with IBEX to visualise components of the OLM in health and disease.

Our refined IBEX protocol permits the use of multiple unconjugated primary antibodies from the same host species (e.g., rabbit polyclonals) over different cycles, unlocking the use of rare niche antibodies needed for specialised tissues such as the retina. We provide a resource of validated antibodies covering the major cell types of the human and mouse retina, many using identical antibody clones for both species (**Supplementary Figure S4**). This work is not intended as a comprehensive or static reference atlas but is intended to evolve by linking into open science initiatives such as the IBEX Imaging Knowledge Base, where further validated antibody markers can be deposited by the research community.[22]

Our spatial proteomic atlas supplements published transcriptomic datasets. There were some marked species-dependent differences within the transcriptome, that can be validated with protein staining, such as the relatively elevated expression of vimentin in horizontal cells in the mouse retina. Vimentin is conventionally considered to be a canonical marker for Müller glia cells. Shaw & Weber however demonstrated horizontal cell expression was different in the mouse retina.[23] Our atlas confirmed this at a protein level, providing direct spatial context and cross-validation of differences between the two species (**Supplementary Figure S5**).

Desmin, for example, had little observed expression in human retinal transcriptomic datasets but was present at a protein level. Comparatively, its expression was found in pericytes and endothelial cells in the mouse retina. This difference may be due to tissue dissociation limitations for scRNAseq[13] or the absence of active transcription of desmin after development. Similarly, rhodopsin transcripts have also been described as problematic in rod-rich retinal samples in scRNAseq with reported expression in all cell types.[13] This was not consistent with protein staining in both mouse and human retina, where rhodopsin was restricted to the outer segments as expected. It was previously reported that rod transcripts can confound cell clustering in scRNAseq due to their abundance and potential cross-contamination during single cell encapsulation.[13]

We identified that the cell adhesion molecule CD44 forms “fibrillar-like” structures at the OLM, colocalising with the F-actin-rich adherens belt (Phalloidin). Identifying all the interacting components of the OLM was challenging with conventional fluorescence confocal microscopy. Therefore, we implemented IBEX in combination with super-resolution microscopy via deconvolution. We were able to observe a novel feature of the OLM in human retina in the form of a structural ‘gap’ at the endfeet of the Müller glia. Although the GS-Phalloidin-CD44 complex was also seen in the mouse retina, this gap was not observed. It may reflect a species-specific structural difference or could be due to the smaller size of mouse retinal structures, which may be at the limit of detection for our imaging approach.

CD44 has been closely linked to the apical processes of the Müller glia and the extracellular matrix.[14, 16] and involved in the development of the retina[17], although its role in the adult human retina is not widely understood. However, CD44 was found to be significantly upregulated in inherited retinal diseases such as retinitis pigmentosa (RP) and in mouse models of RP.[18, 24] Ayten et al. showed that CD44^-/-^ knockout in RP *Pde6b*^*-/-*^ mouse model led to significantly faster photoreceptor degeneration and decreased retinal function.[18] Enrichment of CD44 at the OLM and its close association with the actin cytoskeleton suggest it may play a key role in maintaining the mechanical linkage and signalling between Müller glia and photoreceptors. We directly observe the disruption to the CD44/Phalloidin complex and rosette formation in the *Crb1*^*rd8*^ mouse model as well as the presence of the OLM incorporating photoreceptor outer segment components within the rosettes. This equally suggests that the origin of rosettes forms as an invagination outside the OLM engulfing outer segment components instead of forming within the ONL.[21]

The study has some limitations. Dataset generation was based on post-mortem tissue samples which are static and cannot capture the dynamic changes in cellular processes, although spatial interactions can be documented. There is greater control regarding fixation in mouse tissues, but human retina were often fixed with a post-mortem delay of at least 20 hours. The number of human donor samples is always limited and heterogenous so larger future cohorts will be required to validate our findings. Additionally, any method based on immunohistochemistry has potential issues with antibody specificity and off-target binding, particularly in human-tissues, where antibodies are difficult to validate. This is another reason our study included a mouse retina atlas, in addition to being a good reference for mouse models of disease, as cross species-validation using identical clones adds cell specific assurance. Mouse model knockouts could be used for precise protein validation to confirm their binding in human samples by proxy.

In summary, this research demonstrates the potential of spatial proteomic mapping for the study of the retina. By establishing normative datasets and validated antibody panels, the technology can be applied to other retinal diseases, such as age-related macular degeneration and diabetic retinopathy to reveal new pathological insights. Further technical developments to combine spatial proteomics with other modalities such as spatial transcriptomics, will enable us to construct integrated multi-dimensional atlases of the retina in health and disease to advance biological understanding and impact ophthalmic clinical practice.

## 4 Methods

### 4.1 Human and Mouse Tissue Acquisition and Ethics Approval

Post-mortem human retinal tissue for this study was obtained through Moorfields Biobank (NHS REC approval: 20/SW/0031) with informed consent from donors and eye retrieval provided by NHS Blood and Transplant (NHSBT). C57BL/6J mice were used as the wildtype control. Crb1^*rd8*^/J mouse line was obtained from The Jackson Laboratory (Strain no.: 003392). All animal procedures were performed under a UK Home Office project license (PP1506797) and adhered to the ARVO Statement for the Use of Animals in Ophthalmic and Vision Research.

### 4.2 Tissue Cryopreservation and Sectioning

Human tissues were fixed with 4% PFA for 24 hours at the NHSBT before being exchanged into PBS. Whole eye globes, without the cornea, were shipped to UCL. Crb1^*rd8*^ mice were culled after 28 weeks. Mice underwent cardiac perfusion with 2% paraformaldehyde before their eyes were removed. Both species of retinal tissues were fixed overnight at 4°C in a 1:4 dilution of BD Cytofix (BD Biosciences). Subsequently, tissue samples were cryoprotected by sequential dehydration in 15% and 30% sucrose solutions. Dehydrated tissues were embedded in O.C.T. compound (Cell Path), snap-frozen on dry ice, and stored long-term at -70°C. To ensure strong adhesion of tissue sections during the multiple washing and bleaching cycles, SuperFrost® Plus microscope slides (VWR) were coated with chrome alum gelatin. Frozen tissue blocks were sectioned at 15 μm thickness using a Leica CM1950 cryostat and mounted on the coated slides.

### 4.3 Optimised IBEX Procedure: Immunolabeling, Imaging, and Signal Clearing

Our iterative multiplexed immunohistochemistry technique is based on the IBEX method developed by Radtke et al. with optimizations.[11] The brief workflow is as follows:

Standard IHC blocking protocol was used. Antibodies were diluted according to their respective validated dilutions (see **Supplementary Table S1** for human, and **Supplementary Table S2** for mouse) in a base of PBS + 0.1% Tween (Merck Sigma, Gillingham, UK, P9416). The antibody mix was subsequently applied to the samples, and these were left overnight in a humidified chamber at 4°C.

Most primary antibodies were acquired commercially conjugated. Some acquired purified primary antibodies were conjugated with Antibody Labeling Kits from ThermoFisher, or secondary antibodies were still used for purified primary antibodies only. Secondaries antibodies (see **Supplementary Table S3**) were diluted 1:1000 in PBS and applied to the slides for 1hr at RT in a darkened humidified chamber, limiting ambient light.

### 4.4 Confocal Microscopy

Leica TCS SP8 HyVolution (Leica Microsystems) confocal microscope with 40x objective was used to image mouse and human samples. Super-resolution imaging of the outer limiting membrane was imaged on a Leica Stellaris 8 microscope using a 63x objective. When imaging with the 63x objective, a 6x optical zoom was applied for mouse section imaging and a 5x optical zoom for human section imaging. Leica LIGHTNING deconvolution mode was used when imaging the outer limiting membrane to preserve optimal resolution and signal to noise ratio while imaging in high magnification.

### 4.5 Bleaching and Antibody Removal

Slides were placed into PBS-0.1%-Tween for coverslip removal. Depending on the panel design, fluorophores were either chemically inactivated, or VectaPlex™ Antibody Removal Kit (Vector Laboratories, VRK-1000) was used. Chemical inactivation was carried out using a 1mg/ml solution of Lithium Borohydride (LiBH_4_ Strem Chemicals, UK, 93-0397) in deionised water. The bleaching solution was applied to the tissue 2x 10 minutes under bright ambient room light and washed 3x 10 mins with PBS.

### 4.6 Image Registration and Analysis

Images from each cycle were processed using Imaris software (version 10.2, Imaris Software, Bitplane, Oxford Instruments, Abingdon, UK) for preliminary processing, such as channel naming and Gaussian filtering. Subsequently, an open-source registration software based on SimpleITK was used to precisely align all imaging cycles into a single coordinate system using DAPI or Hoechst nuclear stains as fiducial markers, generating a single multi-channel image file.[25] Quantitative analysis (e.g., fluorescence intensity measurements) was performed using ImageJ/Fiji software or Imaris.

### 4.7 Single-Cell RNA sequencing data

The scRNAseq datasets were obtained from Wang et al. (2022)[26] (Human) and Li et al. (2024)[27] (Mouse) and visualised using Broad Institute Single Cell Portal. Acquired mean expression and percentage expression for specific genes was obtained and reformatted using R^*^ studio.

### 4.8 Statistical Analysis

All statistical analyses were performed using GraphPad Prism (version 9.0.0, GraphPad Software, San Diego, CA, USA). For comparisons between two independent groups, the unpaired, two-tailed unpaired Welch’s t-test was used for data that passed the normality test. Data was presented as the mean ± standard error of the mean (± S.E.M.) A *p* value of less than 0.05 was considered statistically significant. Significance levels in figures were denoted as follows: ^*^p *<* 0.05, ^**^p*<*0.01, ^***^p*<*0.001.

## Supplementary information

Files are accessible to download from: https://doi.org/10.5281/zenodo.17975750

## Acknowledgements

We thank the Institute of Ophthalmology Imaging Unit and BSU for facility access and their technical support. We are grateful to Andrea Radtke and Ryan MacDonald for their helpful discussions and advice. Figures and schematics were created with Biorender.com.

## Author Contribution

Y.M. J.K and C.J.C. conceived and designed the study. Y.M., J.K., Z.D, L.F., R.A., and B.C. performed the experiments. Y.M., J.K and J.H. conducted image processing and data analysis. Z.D, J.H., L.L., C.L., L.F., R.A., Y.T., and B.C. provided technical support and reagents. Y.M. J.K and C.J.C. wrote the manuscript. All authors reviewed and approved the final manuscript.

## Commercial Relationships Disclosure

No relevant conflicts of interests.

