## Supplemental Table 1 for "Spatial proteomic mapping of the human and mouse retina using IBEX"

**Table S1** Primary antibodies and lectins used in immunohistochemistry for fixed-frozen human retina

| Marker | Clone | Conjugate | Isotype | Vendor | Cat. Number | Dilution | RRID |
|---|---|---|---|---|---|---|---|
| DAPI | - | UV | N/A | Merck-Sigma | D9542-1MG | 1:1000 | - |
| GS | GS-6 | AF532 | Mouse IgG2a,$\kappa$ | Merck-Sigma | MAB302 | 1:100 | AB_2110656 |
| Iba1 | - | Purified | Goat IgG | Abcam | Ab5076 | 1:200 | AB_2224402 |
| aSMA | IA4 | AF532 | Mouse IgG2a,$\kappa$ | ThermoFisher | 14-9760-82 | 1:100 | AB_2572996 |
| Desmin | Y66 | AF488 | Rabbit IgG | Abcam | Ab185033 | 1:200 | AB_2892748 |
| Calbindin | - | AF532 | Rabbit Polyclonal | Merck-Sigma | ABN2192 | 1:50 | AB_2935805 |
| PKCa | H-7 | AF680 | Mouse IgG1,$\kappa$ | Santa-Cruz | SC-8393 | 1:100 | AB_628142 |
| ChAT | - | Purified | Goat Polyclonal | Merck-Sigma | AB144P | 1:100 | AB_2079751 |
| B3-Tubulin | TUJ1 | AF488 | Mouse IgG2a,$\kappa$ | Biolegend | 801203 | 1:200 | AB_2564757 |
| Chx10 | E-12 | AF546 | Mouse IgG2a,$\kappa$ | Santa-Cruz | SC-365519 | 1:100 | AB_10842442 |
| GFAP | SMI25 | AF488 | Mouse IgG2b,$\kappa$ | Biolegend | 837508 | 1:200 | AB_2734610 |
| Vimentin | O91D3 | AF555 | Mouse IgG2a,$\kappa$ | Biolegend | 677802 | 1:200 | AB_2565982 |
| CD44 | IM7 | i594 | Rat IgG2b,$\kappa$ | AAT Bioquest | 356128 | 1:100 | AB_2935685 |
| HSP60 | - | Purified | Rabbit Polyclonal | Abcam | ab46798 | 1:200 | AB_881444 |
| Arrestin C | 7G6 | AF488 | Mouse IgG1,$\kappa$ | Merck-Sigma | MABN2636 | 1:200 | AB_2935804 |
| S-Opsin | - | AF532 | Rabbit Polyclonal | Merck-Sigma | AB5407 | 1:100 | AB_177457 |
| STING | E9X7F | AF647 | Rabbit IgG | CST | 90173 | 1:100 | AB_3675228 |
| S100beta | EP1576Y | AF488 | Rabbit IgG | Abcam | Ab196442 | 1:200 | AB_2722596 |
| YH2AX | - | AF546 | Rabbit Polyclonal | ProteinTech | 10856-1-AP | 1:200 | AB_2114985 |
| M/Lopsin | - | i594 | Rabbit Polyclonal | Merck-Sigma | Ab5405 | 1:200 | AB_177456 |
| COL IV | 1042 | e660 | Mouse IgG2b,$\kappa$ | ThermoFisher | 50-9871-82 | 1:200 | AB_2574404 |
| PNA | - | AF488 | N/A | ThermoFisher | L32458 | 1:200 | - |
| Rhodopsin | 1D4 | AF546 | Mouse IgG1,$\kappa$ | Santa-Cruz | SC-57432 | 1:200 | AB_785511 |
