## Supplemental Table 2 for "Spatial proteomic mapping of the human and mouse retina using IBEX"

**Table S2** Primary antibodies and lectins used in immunohistochemistry for fixed-frozen mouse retina

| Marker | Clone | Conjugate | Isotype | Vendor | Cat. Number | Dilution | RRID |
|---|---|---|---|---|---|---|---|
| DAPI | - | UV | N/A | Merck-Sigma | D9542-1MG | 1:1000 | - |
| Desmin | Y66 | AF488 | Rabbit IgG | Abcam | Ab185033 | 1:200 | AB_2892748 |
| S-Opsin | - | AF532 | Rabbit Polyclonal | Merck-Sigma | AB5407 | 1:100 | AB_177457 |
| Chx10 | E-12 | AF546 | Mouse IgG2a,$\kappa$ | Santa-Cruz | SC-365519 | 1:100 | AB_10842442 |
| Hexokinase | C35C4 | i594 | Rabbit IgG | CST | 2024S | 1:100 | AB_2116996 |
| STING | E9X7F | AF647 | Rabbit IgG | CST | 90173 | 1:100 | AB_3675228 |
| CD45 | 30-F11 | AF700 | Rat IgG2b,$\kappa$ | Biolegend | 103128 | 1:100 | AB_493715 |
| S100beta | EP1576Y | AF488 | Rabbit IgG | Abcam | Ab196442 | 1:200 | AB_2722596 |
| Calbindin | - | AF532 | Rabbit Polyclonal | Merck-Sigma | ABN2192 | 1:100 | AB_2935805 |
| HSP60 | - | Purified | Rabbit Polyclonal | Abcam | ab46798 | 1:200 | AB_881444 |
| M/Lopsin | - | i594 | Rabbit Polyclonal | Merck-Sigma | Ab5405 | 1:200 | AB_177456 |
| GFAP | SMI25 | AF647 | Mouse IgG2b,$\kappa$ | Biolegend | 837511 | 1:200 | AB_2734610 |
| PKCa | H-7 | AF680 | Mouse IgG1,$\kappa$ | Santa-Cruz | SC-8393 | 1:100 | AB_628142 |
| GS | GS-6 | AF532 | Mouse IgG2a,$\kappa$ | Merck-Sigma | MAB302 | 1:100 | AB_2110656 |
| Iba1 | - | Purified | Goat IgG | Abcam | Ab5076 | 1:200 | AB_2224402 |
| CD44 | IM7 | i594 | Rat IgG2b,$\kappa$ | AAT Bioquest | 356128 | 1:100 | AB_2935685 |
| B3-Tubulin | TUJ1 | AF488 | Mouse IgG2a,$\kappa$ | Biolegend | 801203 | 1:200 | AB_2564757 |
| P53 | 7F5 | Purified | Rabbit IgG | CST | 2527S | 1:100 | AB_10695803 |
| PNA | - | AF488 | N/A | ThermoFisher | L32458 | 1:200 | - |
| aSMA | IA4 | AF532 | Mouse IgG2a,$\kappa$ | ThermoFisher | 14-9760-82 | 1:100 | AB_2572996 |
| p16 | F-12 | AF546 | Mouse IgG2a,$\kappa$ | Santa-Cruz | SC-1661 | 1:200 | AB_628067 |
| Isolectin B4 | - | AF594 | N/A | ThermoFisher | I21413 | 1:200 | - |
| Calretinin | DAK-Calret1 | AF647 | Mouse IgG1,$\kappa$ | Dako Agilent | M724529-2 | 1:200 | AB_2068519 |
| COL IV | - | Purified | Rabbit Polyclonal | Abcam | ab19808 | 1:100 | AB_445160 |
| Rhodopsin | 1D4 | AF546 | Mouse IgG1,$\kappa$ | Santa-Cruz | SC-57432 | 1:200 | AB_785511 |
| Vimentin | W16220A | AF647 | Rat IgG2a,$\kappa$ | Biolegend | 699308 | 1:200 | AB_2888890 |

**Table S3 Secondary Antibodies used for immunohistochemistry**

| Marker | Clone | Conjugate | Isotype | Vendor | Cat. Number | Dilution | RRID |
|---|---|---|---|---|---|---|---|
| anti-Rabbit IgG | - | AF488 | Donkey IgG | ThermoFisher | A21206 | 1:1000 | AB_2535792 |
| anti-Rabbit IgG | - | AF594 | Donkey IgG | ThermoFisher | A32754 | 1:1000 | AB_2762827 |
| anti-Rabbit IgG | - | AF680 | Donkey IgG | ThermoFisher | A32802 | 1:1000 | AB_2762836 |
| anti-Rat IgG | - | AF594 | Donkey IgG | ThermoFisher | A21209 | 1:1000 | AB_2535795 |
| anti-Goat IgG | - | AF555 | Donkey IgG | ThermoFisher | A32816 | 1:1000 | AB_2762839 |
