## Supplemental Figures for "Spatial proteomic mapping of the human and mouse retina using IBEX"

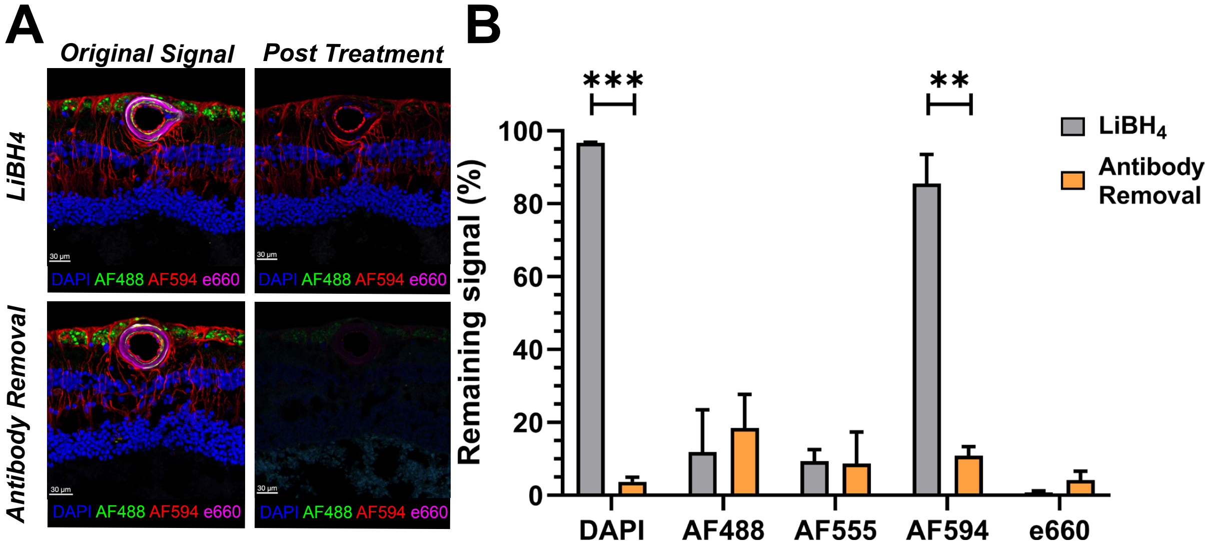


**Figure S1 Antibody removal reagent significantly diminishes fluorescence signal from DAPI and AF594 which is resistant to $LiBH_4$ bleaching. (A)** DAPI or AF594 is not bleached by $LiBH_4$ treatment. Antibody removal reagent attenuates their fluorescence. Scale Bars: 30µm **(B)** Percentage of fluorophore signal remaining after 20 minutes of $LiBH_4$ or antibody removal reagent treatment. Mean ± SEM, n=3 different human retinas. DAPI and AF594 signal is significantly attenuated by antibody removal compared to $LiBH_4$ (T-test: **p<0.01, ***p<0.001)

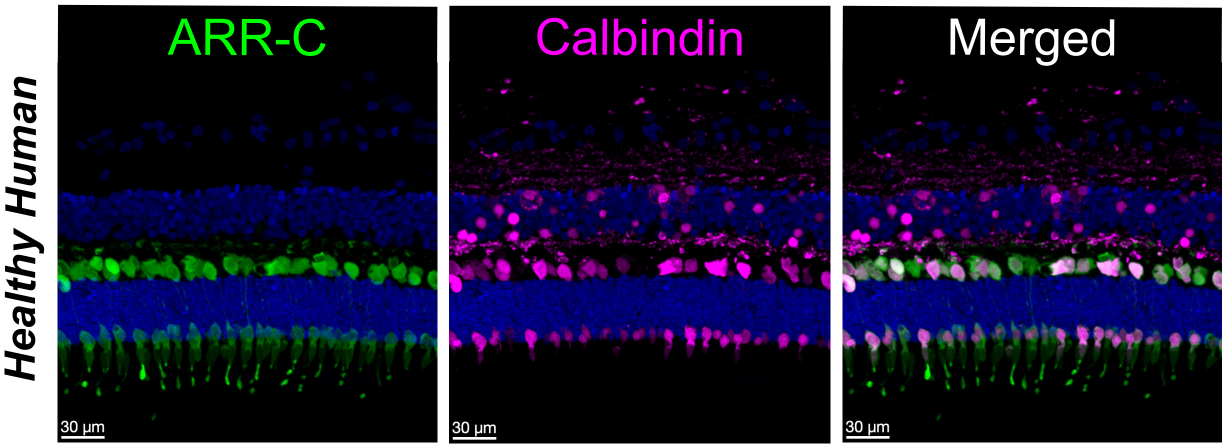


**Figure S2** Composite image generated using IBEX demonstrates ARR-C staining to illustrate cones and calbindin within the inner segment region of a healthy human central retina. Calbindin expression is observed within the inner segments of ARR-C+ cones.

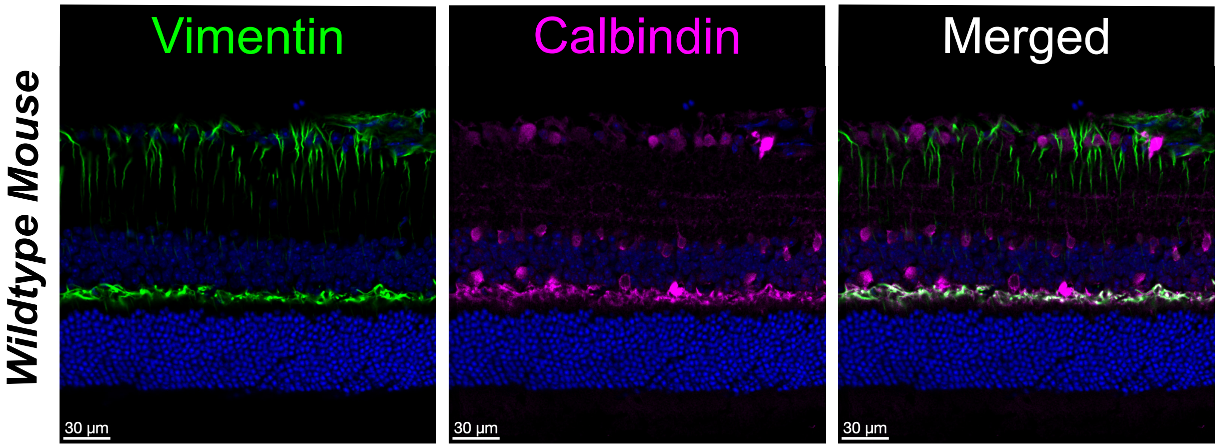


**Figure S3** Composite image generated using IBEX shows Vimentin and Calbindin staining in a wildtype mouse retina. Vimentin expression in mouse retina at the OPL region colocalises with calbindin.

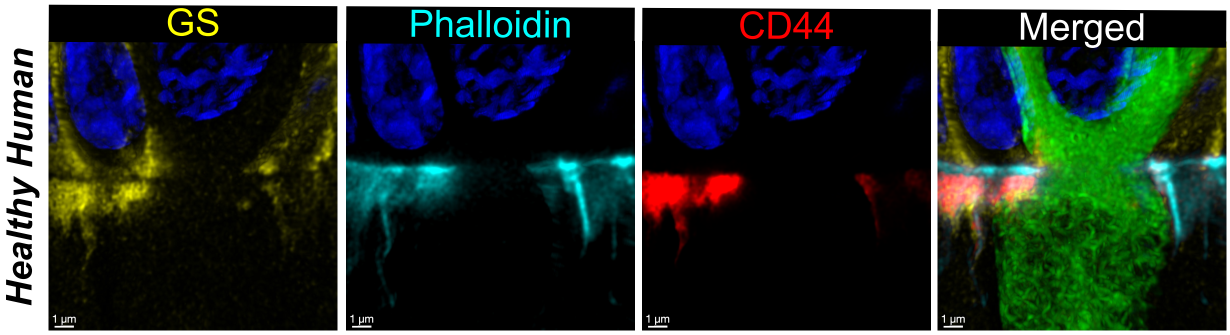


**Figure S4** Super-resolution imaging of the GS-Phalloidin-CD44 complex in the human retina. A linear gap can be observed at the endfeet of Müller Glia colocalising with the F-actin-rich adherens belt (Phalloidin)

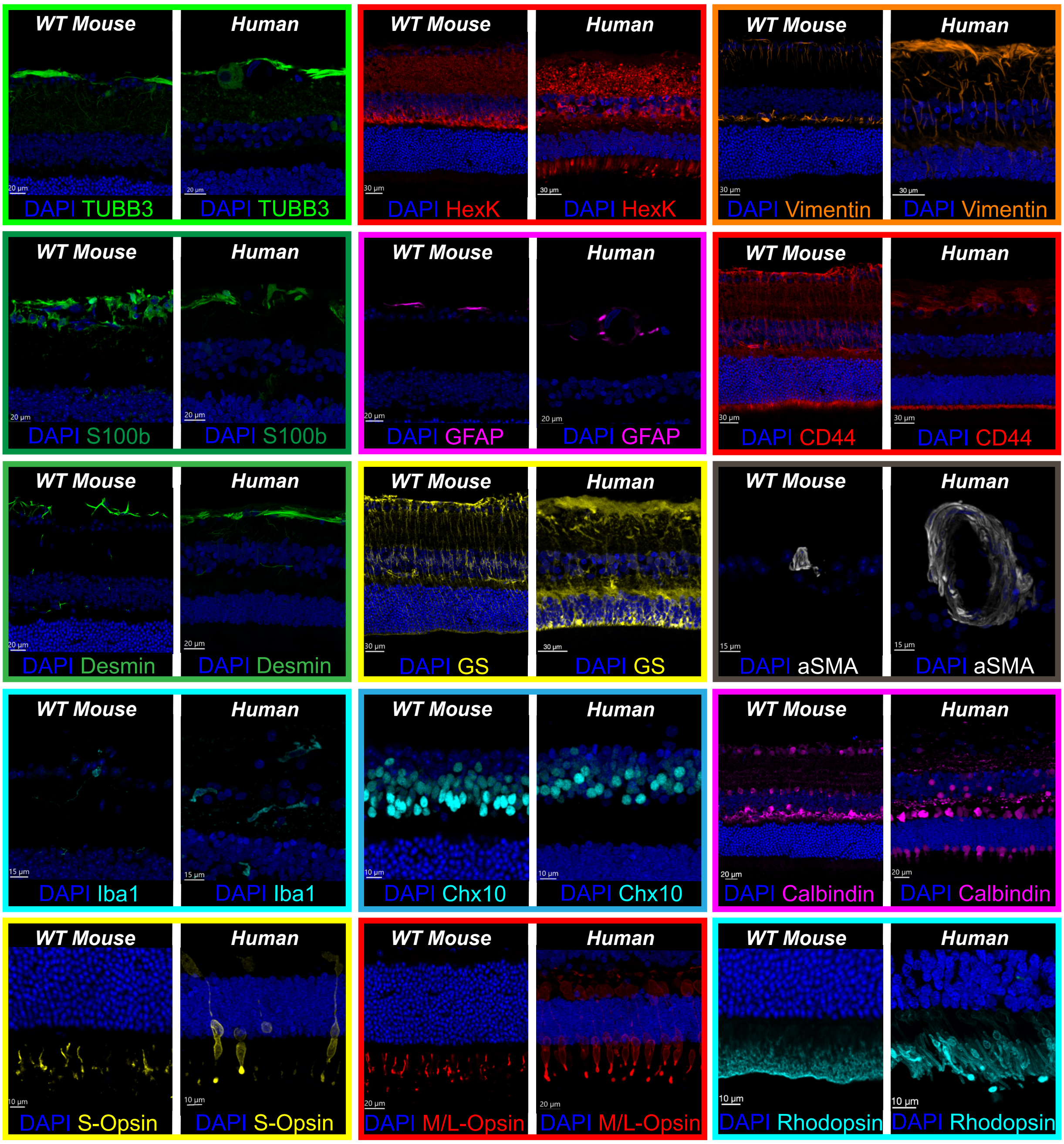


**Figure S5** Cross-reactive protein marker comparisons in wildtype mouse and healthy human retinas. All images of the fifteen cross-reactive markers were generated using IBEX.
